# Allele frequency spectra in structured populations: Novel-allele probabilities under the labelled coalescent

**DOI:** 10.1101/2019.12.20.883629

**Authors:** Marcy K. Uyenoyama, Naoki Takebayashi, Seiji Kumagai

**Author notes:** Corresponding author: Marcy K. Uyenoyama, Department of Biology, Box 90338, Duke University, Durham, NC 27708-0338, USA.

## Abstract

We address the effect of population structure on key properties of the Ewens sampling formula. We use our previously-introduced inductive method for determining exact allele frequency spectrum (AFS) probabilities under the infinite-allele model of mutation and population structure for samples of arbitrary size. Fundamental to the sampling distribution is the novel-allele probability, the probability that given the pattern of variation in the present sample, the next gene sampled belongs to an as-yet-unobserved allelic class. Unlike the case for panmictic populations, the novel-allele probability depends on the AFS of the present sample. We derive a recursion that directly provides the marginal novel-allele probability across AFSs, obviating the need first to determine the probability of each AFS. Our explorations suggest that the marginal novel-allele probability tends to be greater for initial samples comprising fewer alleles and for sampling configurations in which the next-observed gene derives from a deme different from that of the majority of the present sample. Comparison to the efficient importance sampling proposals developed by De Iorio and Griffiths and colleagues indicates that their approximation for the novel-allele probability generally agrees with the true marginal, although it may tend to overestimate the marginal in cases in which the novel-allele probability is high and migration rates are low.

## 1 Introduction

Denote the allele frequency spectrum (AFS) of a sample of *n* genes at a single, non-recombining locus by

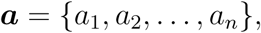

in which *a*_*i*_ represents the number of alleles observed in multiplicity *i*. Under the infinite-alleles model of mutation, for which each mutational event generates a novel allelic class, the Ewens Sampling Formula (ESF, Ewens 1972) provides the probability of AFS ***a*** in the absence of population structure:

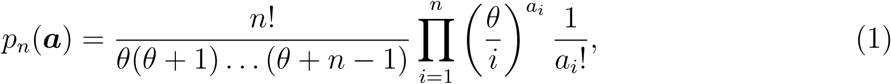

for *p*_*n*_(***a***) = Pr(***a***|**Φ**) with **Φ** = {*θ*}, the scaled rate of mutation,

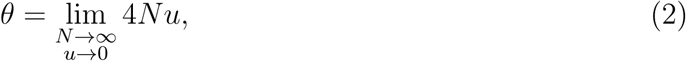

in which *u* represents the per-gene, per-generation rate of mutation and 2*N* the number of genes in the population eligible to leave descendants.

The publication in *Theoretical Population Biology* of the ESF numbers among the watershed moments in evolutionary genetics. Kingman (2000) has recognized this breakthrough as a factor that precipitated the development of coalescence theory. It may still be not widely appreciated that the recursion developed by Karlin and McGregor (1972) to support the ESF is a coalescence argument. Moreover, their argument describes a *labelled* coalescence process, under which the observed pattern of genetic variation provides information about the genealogical history of a sample immediately ancestral to the most recent evolutionary event.

In a remarkable paper, Stephens and Donnelly (2000) elucidated the significance of insights gained from the ESF for the genealogical history of a sample of genes. It unified a number of approaches, notably importance sampling (IS), to the determination of the likelihood of an evolutionary model. In particular, the Griffiths-Tavaré approach (*e.g.*, Griffiths and Tavaré 1994b) to solving recursions in likelihoods corresponds to an IS procedure (Felsenstein *et al.* 1999; Stephens and Donnelly 2000). Under this approach, backward-in-time sequences of ancestral samples culminating in the most recent common ancestor (MRCA) of the sample are proposed using the evolutionary rates of the forward-in-time process. Drawing on properties of the ESF, Stephens and Donnelly (2000) developed a new and more efficient class of IS proposals by connecting the distribution of the immediate ancestor of a sample to the distribution of the next-observed gene conditional on that sample.

Within this class of IS proposals are those developed by De Iorio and Griffiths and colleagues (*e.g.*, De Iorio and Griffiths 2004a,b; De Iorio *et al.* 2005) for general models of mutation in structured populations. Their highly-efficient family of proposals draws on properties of the ESF to approximate the novel-allele probability: the probability the next-observed gene represents a novel allelic class, given the AFS of the present sample.

We have developed an inductive method for determining the probabilities of all possible allele frequency spectra (AFSs) under the infinite-alleles model of mutation for a two-deme population (Uyenoyama *et al.* 2019). While our model constitutes perhaps the simplest within the domain addressed by De Iorio and Griffiths and colleagues, it is the only case beyond the ESF itself for which exact AFS probabilities are known. Here, we compare the actual distribution of the immediate ancestor of a sample to their IS proposals. As this family of proposals, including that of Stephens and Donnelly (2000), is based on characterizing the distribution of the next-sampled gene, we address in particular the probability that the next-sampled gene represents a novel allelic class.

Key to the IS proposals is the approximation that the novel-allele probability is independent of the AFS of the present sample. Our previous study (Uyenoyama *et al.* 2019) showed that this signature property of the ESF is not preserved under population structure. An unanticipated finding of our present analysis is that the approximate novel-allele probability of De Iorio and Griffiths and colleagues is in fact similar to the marginal novel-allele probability, the mean novel-allele probability over all possible AFSs of the sample to which the last gene is added.

We begin with a review of a number of coalescence-based methods that exploit the information contained in the pattern of genetic variation observed in a sample of genes to characterize the ancestor of the sample. We then enumerate some key properties of the ESF for unstructured populations and address the extent to which those properties are preserved under population structure. For a two-deme population under infinite-alleles mutation, we explore qualitative trends in the novel-allele probability and assess the IS proposals of De Iorio and Griffiths (2004b). We present an inductive method for determining the marginal novel-allele probability for samples of arbitrary size, noting that the IS proposal for the novel-allele probability of De Iorio and Griffiths and colleagues approximates this marginal probability.

## 2 Labelled histories

### 2.1 Genealogical histories conditional on observed variants

Observation of the pattern of variation in a sample constrains the domain of possible states of lineages ancestral to the sampled genes. For example, the assumption of low rates of coalescence and mutation excludes the possibility of immediate coalescence between non-identical genes. Beyond such forbidden transitions, the possible ancestral states do not necessarily occur with uniform probabilities.

Wiuf and Donnelly (1999) addressed the genealogical histories implied by the observation of a single mutation. In the absence of homoplasy, all lineages bearing the mutation must coalesce with one another more recently than any coalesce with other lineages. Accordingly, the observed pattern of variation implies a topology that includes a branch from which all mutation-bearing lineages and no other lineages descend. The Wiuf and Donnelly (1999) method generates topologies by proposing an ancestor state with probability corresponding to the probability that the state has a genealogical history consistent with the pattern of mutation. Because the proposal distribution is in fact the desired probability, it is the optimal proposal distribution. They obtained an exact expression for the probability of the set of topologies on which the observed pattern of variation has positive probability. Wiuf and Donnelly (1999) used their method to address the age of a unique mutation segregating in a sample of genes derived from a panmictic population.

Leman *et al.* (2005) incorporated the Wiuf and Donnelly (1999) approach in developing an IS analysis of genetic variation in a noncoding region assumed to show absolute linkage to the paracentric inversion that contributes to reproductive isolation between a pair of Drosophila species. This study generated maximum likelihood estimates of the time since speciation and the effective population sizes of the extant species and their ancestor species. The observed pattern of genetic variation in the structured sample corresponded to the number of segregating mutations of 7 types, reflecting whether a mutation is absent (*a*) from the subsample derived from a given species, segregating (*s*, present in some but not all lineages in the subsample), or fixed (*f*, present in all lineages in the subsample). Observation of multiple mutations present in the subsample from each species but absent from the subsample from the other species implies an MRCA of each subsample more recent than any cross-species coalescence event (see Fig. 1). Building on the principles introduced by Wiuf and Donnelly (1999), Leman *et al.* (2005, Appendix B) developed a recursion in the probability of a topology consistent with both the mutational array and the process of speciation, including the forward progression from a single ancestor species to complete reproductive isolation at the observed locus. Each IS proposal of the complete genealogical history entails generation of a topology of the full sample and then placement of mutations on the tree. As the proportion of random trees on which the observed data have positive probability is on the order of 7.7×10^−9^, incorporation of the modified Wiuf and Donnelly (1999) approach greatly improves the efficiency of the IS analysis by ensuring that all proposed trees are consistent with the observed data.

**Figure 1:**
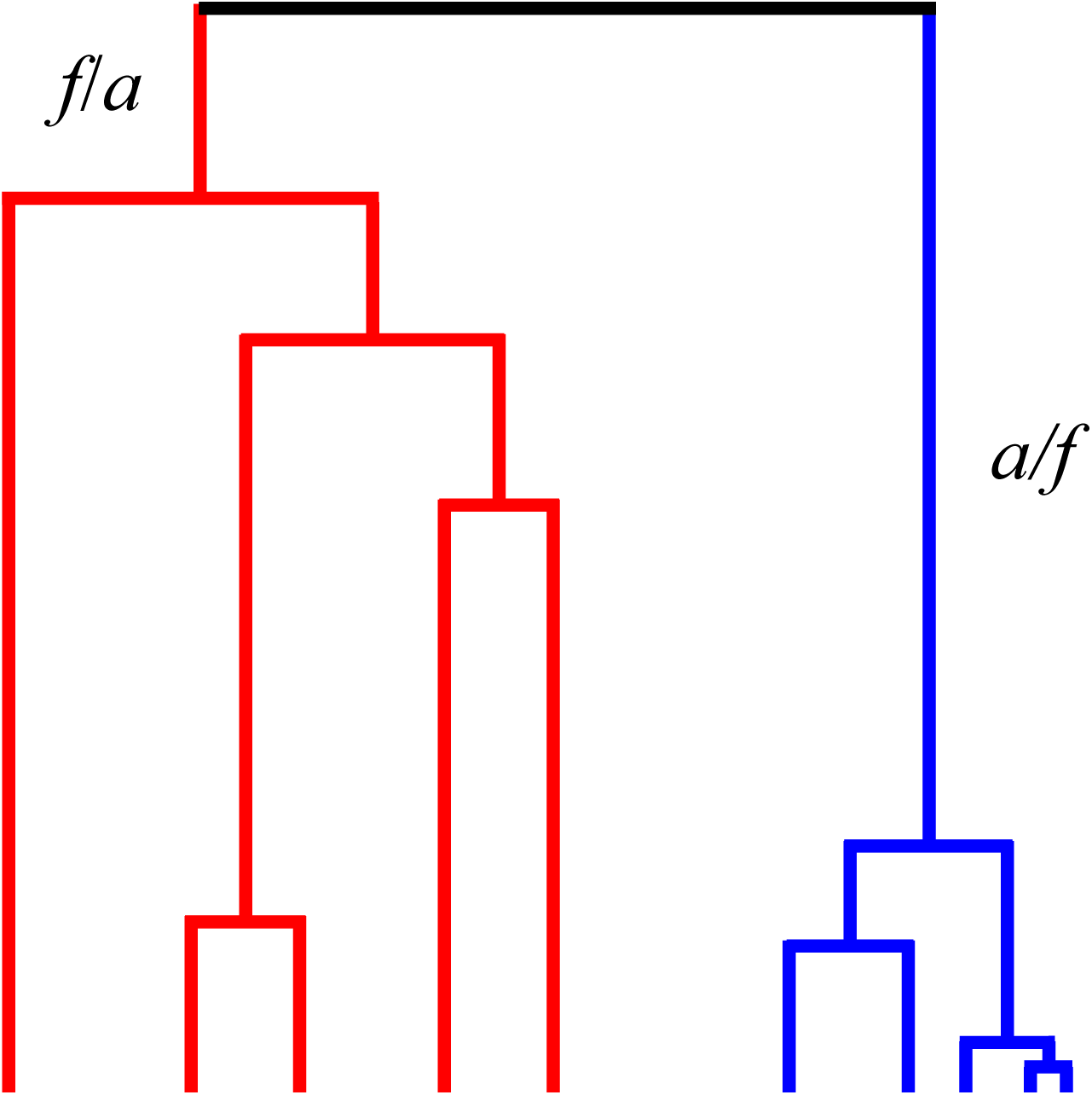
Topology consistent with observation of fixed mutational differences between sub-samples derived from each of two species (red and blue). Mutations arising on the branch labelled *f/a* are fixed (*f*) in the subsample derived from the red species and absent (*a*) from the subsample derived from the blue species. Similarly, mutations arising on the branch labelled *a/f* occur only in the subsample derived from the blue species.

### 2.2 Karlin-McGregor recursion

#### Next-sampled gene

Ewens’s (1972) own derivation of the ESF (1) proceeded from the “remarkable intuitive insight” (Karlin and McGregor 1972) that the probability that the last (*n*^*th*^) gene added to a sample of size *n* − 1 represents a novel allele corresponds to

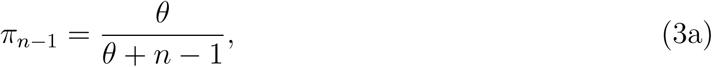

with the probability that the last gene belongs to an allelic class already represented in the sample in multiplicity *i* (*i* = 1, 2, …, *n* − 1) given by

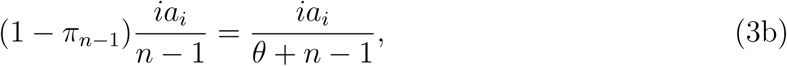

for *a*_*i*_ denoting the number of alleles present in the sample in multiplicity *i* (1).

#### Labelled coalescent argument

To establish the ESF without prior knowledge of the distribution of the allelic type of the next-sampled gene (3), Karlin and McGregor (1972) used a labelled coalescent argument to produce a recursion in the AFS probabilities and showed that the ESF (1) satisfies it.

As in Uyenoyama *et al.* (2019), we denote the present (descendant) sample by *D* = ***a***, for ***a*** the allele frequency spectrum (AFS). In the case of structured populations, for example, ***a*** provides information on the multiplicities and locations of the alleles in the sample. Let *T* represent the single evolutionary event that separates descendant *D* = ***a*** from its immediate ancestor *A* = ***b***. For example, in the model (6) we will address here, the evolutionary event may correspond to migration, mutation, or coalescence. In general, the event may reflect any evolutionary process, including recombination. The likelihood of the model together with its parameters (**Φ**) corresponds to the solution of a recursion over the most recent evolutionary event:

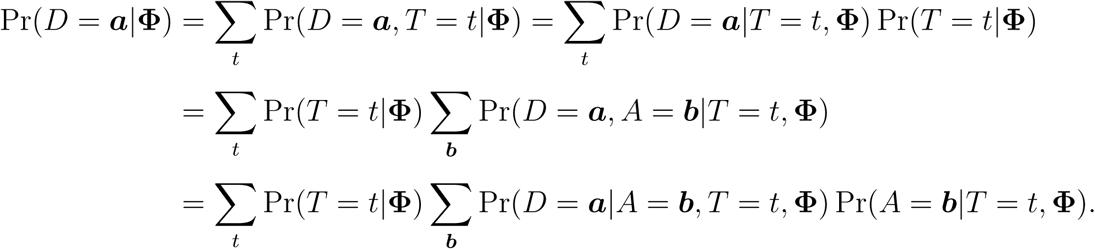

Using that the ancestor state is independent of the next evolutionary event forward in time,

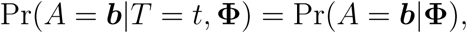

we obtain

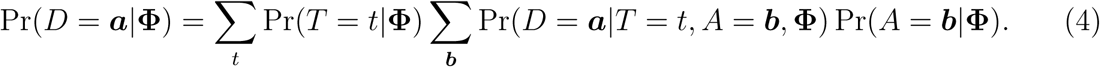

Similar to what has now become a standard coalescence approach, this recursion may be read as conditioning first on the evolutionary event (*T*). The information contained in the observation of the labelling (allelic class) of the sampled genes affects Pr(*D* = ***a***|*T* = *t, A* = ***b*, Φ**), which links the ancestor to the descendant. In many assignments of the ancestor AFS (***b***), this term is likely to be zero. Appendix A presents the recursion under the infinite-alleles model in unstructured populations.

Recursion (4) applies in quite general contexts. Restricted to the infinite-alleles model of mutation, it can be solved inductively, by progressively incrementing sample size, number of distinct allelic classes, and number of singleton alleles, to produce AFS probabilities for samples derived from structured populations (Uyenoyama *et al.* 2019).

#### Conditional probabilities

Karlin and McGregor (1972) observed that ratios of AFS probabilities may be interpreted as conditional probabilities: for example, the probability of an AFS formed upon sampling an additional gene, given the present AFS. In the case of the ESF, (3a) corresponds to

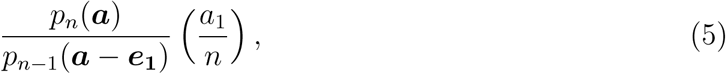

in which the first factor represents the conditional probability of a full sample (size *n*) with AFS ***a*** given that the penultimate sample (size *n* − 1) has AFS ***a*** − ***e***_**1**_, and the second the probability that the last (*n*^*th*^) gene sampled corresponds to one of the *a*_1_ singleton alleles. This interpretation of ratios of AFS probabilities as conditional probabilities is key to the insightful IS proposals developed by De Iorio and Griffiths (2004b) for generalized models of mutation in structured populations. Further, we show in the Appendix B that it provides another means of obtaining the novel-allele probability (3a) directly, even without derivation of the full ESF (1).

### 2.3 Distribution of the ancestor given the descendant in unstructured populations

A number of works have explored methods for approximating the distribution of the immediate ancestor of an observed sample in more general contexts (*e.g.*, Hoppe 1987; Griffiths and Tavaré 1994a,b; Stephens and Donnelly 2000; Tavaré 2004; De Iorio and Griffiths 2004b). By addressing the distribution of the next-sampled gene, Stephens and Donnelly (2000) developed a more efficient class of importance sampling (IS) proposal distributions for generating genealogical histories. Hobolth *et al.* (2008) presented a comparison of the IS proposal distributions of Griffiths and Tavaré (1994a) and Stephens and Donnelly (2000) under the infinite-sites model.

Stephens and Donnelly (2000) noted that determination of the exact distribution of the immediate ancestor of the lineages in a sample conditional on the descendant sample is tantamount to full knowledge of AFS probabilities. As an illustration, we use the full solution provided by the ESF (1) to describe this distribution under the infinite-alleles model of mutation in an unstructured population (Appendix C).

## 3 Structured populations

### 3.1 Key properties of the ESF

We address the extent to which the sampling properties of the ESF are preserved under infinite-alleles mutation in subdivided populations, in which evolutionary events include migration as well as mutation and coalescence. In the two-deme setting explored by Uyenoyama *et al.* (2019), deme *i* (*i* = 0, 1) comprises an effective number of 2*N*_*i*_ genes with backward migration rate *m*_*i*_ and novel alleles arise at the per-gene, per-generation mutation rate of *u*, implying 4 parameters:

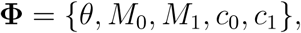

in which

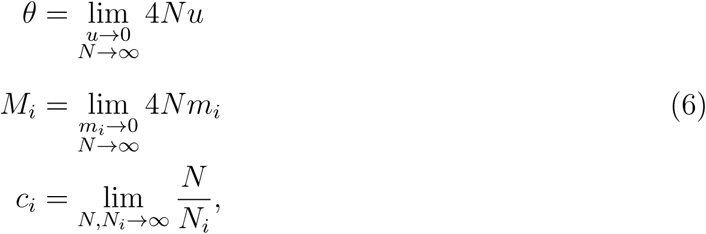

for *N* an arbitrary constant that goes to infinity at a rate comparable to the *N*_*i*_ (compare (1)). AFS ***a*** now comprises elements of the form *a*_*ij*_, corresponding to the number allelic classes that have *i* replicates in the subsample derived from deme 0 and *j* replicates in the subsample derived from deme 1. Similarly, ***e***_***ij***_ denotes a unit vector, with unity in the *ij*^*th*^ position and zeros elsewhere. For clarity, we use 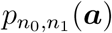 to denote the probability of AFS ***a***, with the subscript explicitly indicating the number of genes (*n*_0_ and *n*_1_) derived from the two demes.

Determining the distribution of the ancestor AFS in general contexts would be facilitated if some key properties of the ESF were universally preserved. As Appendix C illustrates, such properties in the case of structured populations might include that

1. the probability that the next-sampled gene represents a novel allele depends on the parameters of the model and the sampling configuration (demic origin of the genes) but not on the particular AFS observed in the penultimate sample (prior to the addition of the last gene) and
2. for a given most recent evolutionary event (migration, mutation, or coalescence) in a specified deme, all genes in the ultimate sample (after the addition of the last gene) residing in that deme have a uniform probability of having participated in that evolutionary event.

These properties of the ESF do not in fact extend to structured populations (Uyenoyama *et al.* 2019). In particular, the probability of sampling a novel allele depends on AFS of the present sample. Even so, the IS proposals based on these properties appear to be the most efficient available for generalized mutation and population structures (De Iorio and Griffiths 2004b; De Iorio *et al.* 2005). In particular, the IS proposals of De Iorio and Griffiths (2004b) for the probabilities of sampling a gene that represents a novel allele or an already-observed allelic class represent solutions of linear systems of equations that those quantities must satisfy if properties possessed under the ESF framework were preserved under the generalized framework.

### 3.2 De Iorio-Griffiths IS proposals

Given observation of AFS ***a*** in a sample comprising *n*_*i*_ genes derived from deme *i* (*i* = 0, 1), the probability that a gene sampled from deme 0 represents a novel allele corresponds to

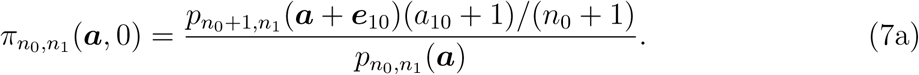

This expression reflects the characterization (5) of Karlin and McGregor (1972) of the last-sampled gene in terms of a conditional probability, with the factor (*a*_10_+1)*/*(*n*_0_+1) denoting the probability that the last-sampled gene is a singleton allele. Similarly, the probability that the last-sampled gene represents an allele already present in the sample in multiplicity *x*_0_ in deme 0 and *x*_1_ in deme 1 corresponds to

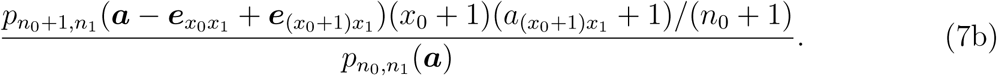

Analogous expressions arise in the case in which the next-sampled gene derives from deme 1.

The IS proposals of De Iorio and Griffiths (2004b) incorporate an approximation to 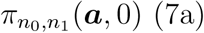, which reduces in the case at hand (6) to

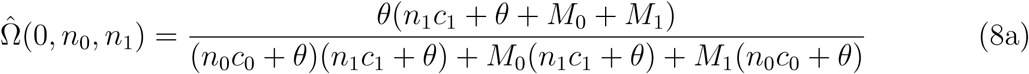

(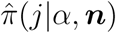 in their notation). Their approximation for the probability (7b) that the last-sampled gene belongs to a particular allelic class for which the sample already contains *x*_0_ replicates in deme 0 and *x*_1_ replicates in deme 1 is

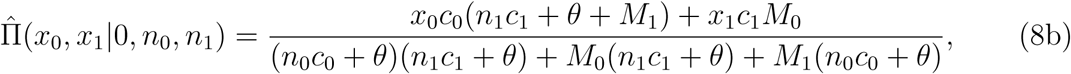

for *x*_0_ > 0 or *x*_1_ > 0.

Similar expressions hold for cases in which the last gene is derived from deme 1:

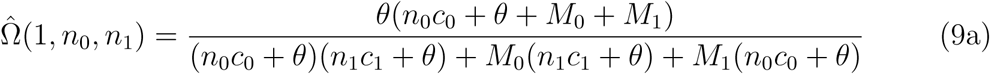

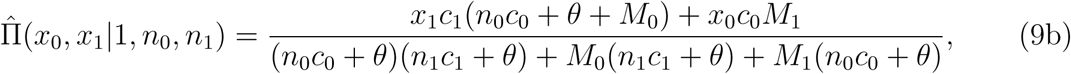

for *x*_0_ > 0 or *x*_1_ > 0.

### 3.3 Marginal probability of a novel allele

For our two-deme model (6), we present an inductive method to determine the probability that a gene sampled from a specified deme and added to a sample comprising *n*_*i*_ genes from deme *i* (*i* = 0, 1) represents a novel allele, marginalized over all possible allele frequency spectra observed in the initial sample. As we can in principle obtain all AFS probabilities, this marginal probability might be obtained from expressions such as (7a), corresponding to sampling of the next gene from deme 0:

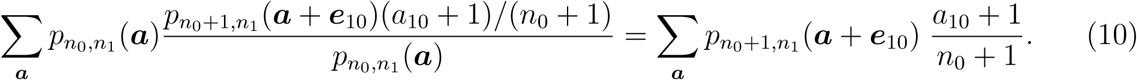

On the left is the conditional probability, expressed as a ratio as in (5), that a gene derived from deme 0 represents a novel allele multiplied by the probability of the penultimate sample (prior to the addition of that gene). Our method, implemented in the supplemental Mathematica notebook accompanying this article, permits determination of the marginal probability of a novel allele directly, without prior knowledge of all AFS probabilities.

Under the infinite-alleles model, we need follow a lineage up to only the most recent mutation or coalescence event (compare Griffiths and Lessard 2005). Regardless of the number of older mutations a lineage may have accumulated, a mutation represents the origin of an allelic class in the population. Level *ℓ* of the full *n*-gene genealogy corresponds to the segment that comprises *ℓ* lineages. That the last-sampled gene represents a novel allele entails either that the focal lineage terminates in a mutation on level *ℓ* (*ℓ* = 3, …, *n*) of the gene genealogy or that the focal lineage persists to level 2 and a mutation occurs in either of the remaining lineages on that level.

On level *ℓ*, state *i* (0 ≤ *i* ≤ *ℓ* − 1) corresponds to the residence of the focal lineage in deme 0 together with *i* non-focal lineages; similarly, state *i* (*ℓ* ≤ *i* ≤ 2*ℓ* − 1) corresponds to the residence of the focal lineage in deme 1 together with *i* − *ℓ* non-focal lineages. From state *i* on level *ℓ*, the total rate of change corresponds to

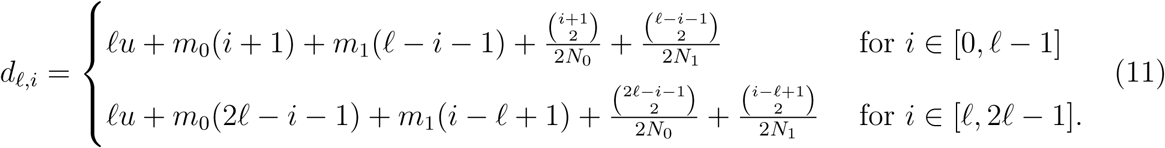

Because a coalescence event involving the focal lineage implies that the focal lineage shares its allelic state with at least one non-focal lineage, we exclude such an event in determining the probability that the focal gene represents a novel allele. Accordingly, the most recent event backward in time may correspond to a transition to a transient state on level *ℓ*, reflecting migration, or to termination of the level, either through coalescence between a pair of non-focal lineages or through mutation in the focal lineage or in a non-focal lineage. Termination of the focal lineage by mutation implies that it represents a novel allele.

To determine the probability that the process terminates on level *ℓ*, with a mutation in the focal lineage, or that it continues on to level *ℓ* − 1, we describe instantaneous rates of within-level and between-level transitions. Matrix ***Û***_***ℓ***_, a square matrix with 2*ℓ* rows and columns, provides the probabilities of transitions through migration, with the *ij*^*th*^ element denoting the probability that the most recent event back in time corresponds to a transition from state *i* to state *j*. For the residence of the focal gene in deme 0 (*i* ∈ [0, *ℓ* − 1]),

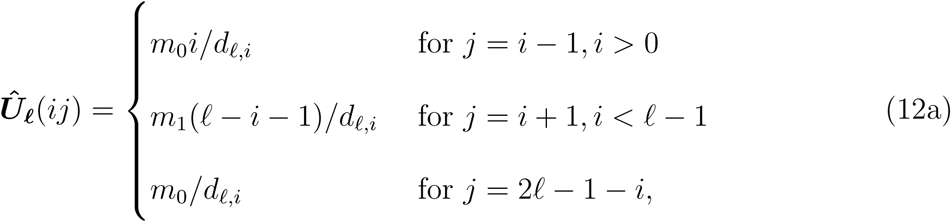

with all other elements set to zero. These transitions respectively denote backward migration of a non-focal lineage presently in deme 0, of a non-focal lineage presently in deme 1, and of the focal lineage itself, with ***Û***_***ℓ***_(*ij*) = 0 for other values of *i* and *j*. Similarly, for the residence of the focal gene in deme 1 (*i* ∈ [*ℓ*, 2*ℓ* − 1]), elements of ***Û***_***ℓ***_ correspond to

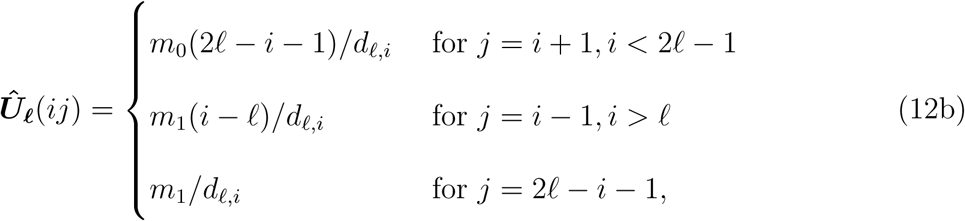

again respectively denoting backward migration of a non-focal lineage presently in deme 0, of a non-focal lineage presently in deme 1, and of the focal lineage itself.

Matrix 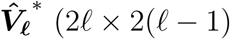, for 2(*ℓ* − 1) the number of states on level *ℓ* − 1), provides the probabilities of between-level transitions that do not involve the focal lineage: a mutation in a non-focal lineage or coalescence between a pair of non-focal lineages. State *i* (*i* ∈ [0, *ℓ* − 1]) denotes the residence of the focal gene in deme 0 together with *i* non-focal lineages, with the remaining (*ℓ* − 1 − *i*) lineages (all non-focal) residing in deme 1. Elements of 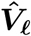 include

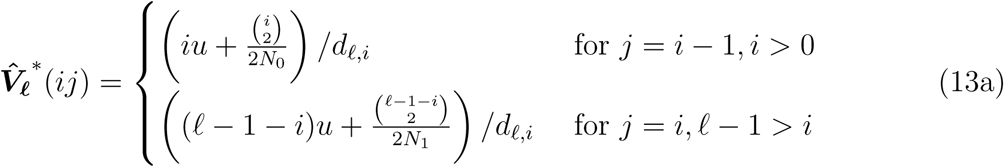

with other entries and unmeaningful expressions (*e.g.*, 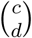 with *c* < *d*) set to zero. Similarly, for the focal gene residing in deme 1 (*i* ∈ [*ℓ*, 2*ℓ* − 1]) together with (*i* − *ℓ*) non-focal lineages and the remaining (2*ℓ* − 1 − *i*) non-focal lineages in deme 0,

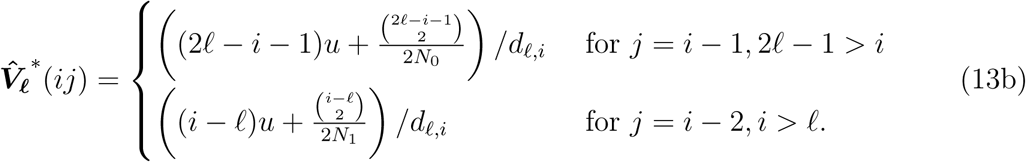

Vector ***T***_***ℓ***_ (2*ℓ* × 1) provides rates of termination with the focal gene representing a novel allele:

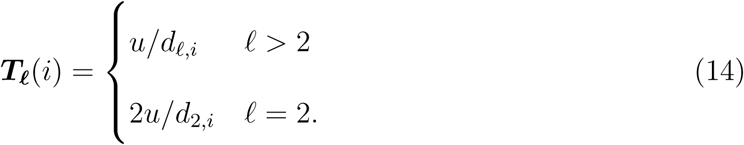

Here, the expression for level *ℓ* > 2 reflects a mutation in the focal lineage and the expression for level *ℓ* = 2 reflects a mutation either in the focal lineage or in the single remaining non-focal lineage.

We now replace the elements of 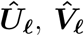, and ***T***_***ℓ***_ by their limiting values as described in (6): for *ℓ* > 2 for example,

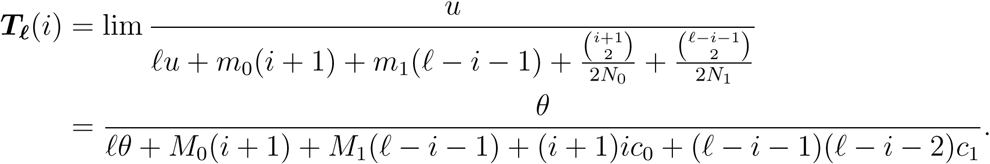

We describe as transient the states among which transitions reflecting migration occur as indicated by ***Û***_***ℓ***_. The columns of 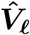 and elements of 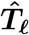 correspond to exit states, indicating termination of level *ℓ*. As shown in Appendix A of Uyenoyama *et al.* (2019), the probability that a process presently in transient state *i* exits through state *j*, representing an event not involving the focal lineage, corresponds to the *ij*^*th*^ element of

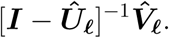

The probability of an exit through mutation in the focal gene is similar but with 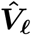 replaced by ***T***_***ℓ***_.

We use ***W***_***n***_ to denote the vector of probabilities that the focal gene, added to a sample of size *n* − 1 is novel. For *i* ∈ [0, *n* − 1], the *i*^*th*^ element, ***W***_***n***_(*i*), provides the probability that a gene, obtained from deme 0 to form an *n*-gene sample comprising *n*_0_ = *i* + 1 genes from deme 0 and the remainder from deme 1, represents a novel allele. Similarly, ***W***_***n***_(*i*) for *i* ∈ [*n*, 2*n* − 1] provides the probability that a gene, obtained from deme 1 to form an *n*-gene sample comprising *n*_1_ = *i* − *n* + 1 genes from deme 1 and the remainder from deme 0, represents a novel allele.

We determine ***W***_***n***_ for a sample of arbitrary size by induction, beginning with *n* = 2, for which

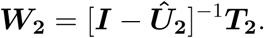

For example, under symmetry between the demes in migration rates and effective numbers,

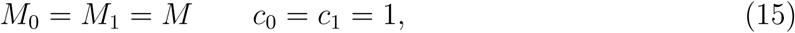

the vector of probabilities reduces to

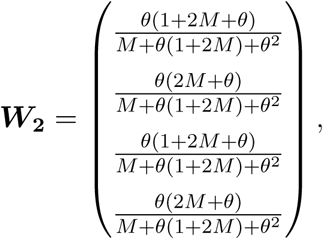

confirming the results of Hudson (1990). For arbitrarily large migration rates (*M* → ∞), every element of ***W***_**2**_ in this case reduces to

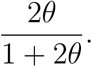

This expression agrees with the expression under the ESF (3a), noting that the total effective number of the combined demes is 2*N*, which implies that *θ* in (3a) corresponds in this case (*M* → ∞) to our 2*θ*.

In general (*ℓ* > 2), a process in state *i* on level *ℓ* may terminate immediately, with a mutation in the focal lineage, with probability given by the *i*^*th*^ element of

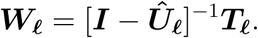

Otherwise, the process proceeds to level *ℓ* − 1, with the probability that the focal gene represents a novel allele given by

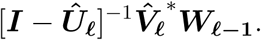

Accordingly, ***W***_***n***_ may be determined inductively, from

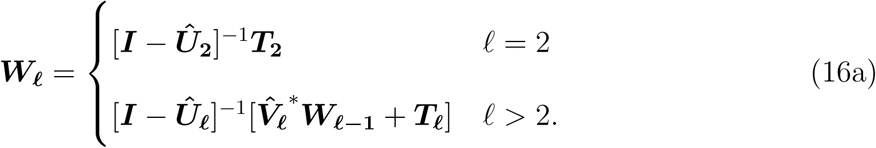

For arbitrary sample size *n* and *z* the level on which the mutation that establishes the focal gene as a new allele occurs,

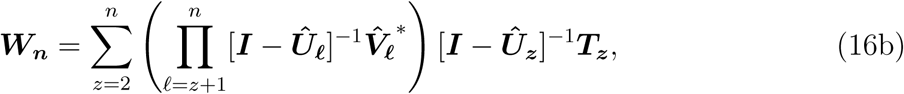

in which the matrix product begins on the left with 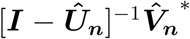 and ends on the right with 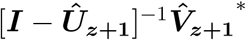.

In panmictic populations, the probability that the lineage of the last-sampled gene terminates in a mutation on a given genealogical level is uniform across levels (Appendix A of Redelings *et al.* 2015). This property is not preserved under population subdivision.

## 4 Assessment of IS proposals

We assess characteristics of the IS proposals given in Section 3.2 in a two-deme setting (6).

### 4.1 Next-observed gene

We address the probability that the next gene sampled from a specified deme represents a novel allele. We compare the IS proposal (8a) of De Iorio and Griffiths (2004b) to the exact marginal novel-allele probability (16).

A total of 2193 AFSs are possible for a sample size *n* = 10, including all possible sample configurations (assignments of *n*_0_ and *n*_1_ for *n*_0_ + *n*_1_ = 10). For 10-gene samples derived wholly from a single deme (*n*_0_ = 10 or *n*_1_ = 10), the number of AFSs corresponds to 42, the number under the ESF (1) and the answer to the ultimate question of life, the universe, and everything. Figure 2 presents the 42 AFSs for a penultimate sample of size *n* = 10, all derived from deme 1, ranked by the probability that an additional (11^*th*^) gene, sampled from deme 0, represents a novel allele. The blue horizontal line corresponds to the marginal novel-allele probability across AFSs, with AFS weighted by its probability, obtained recursively using (16). The horizontal black line, corresponding to the novel-allele probability expected under the ESF

**Figure 2:**
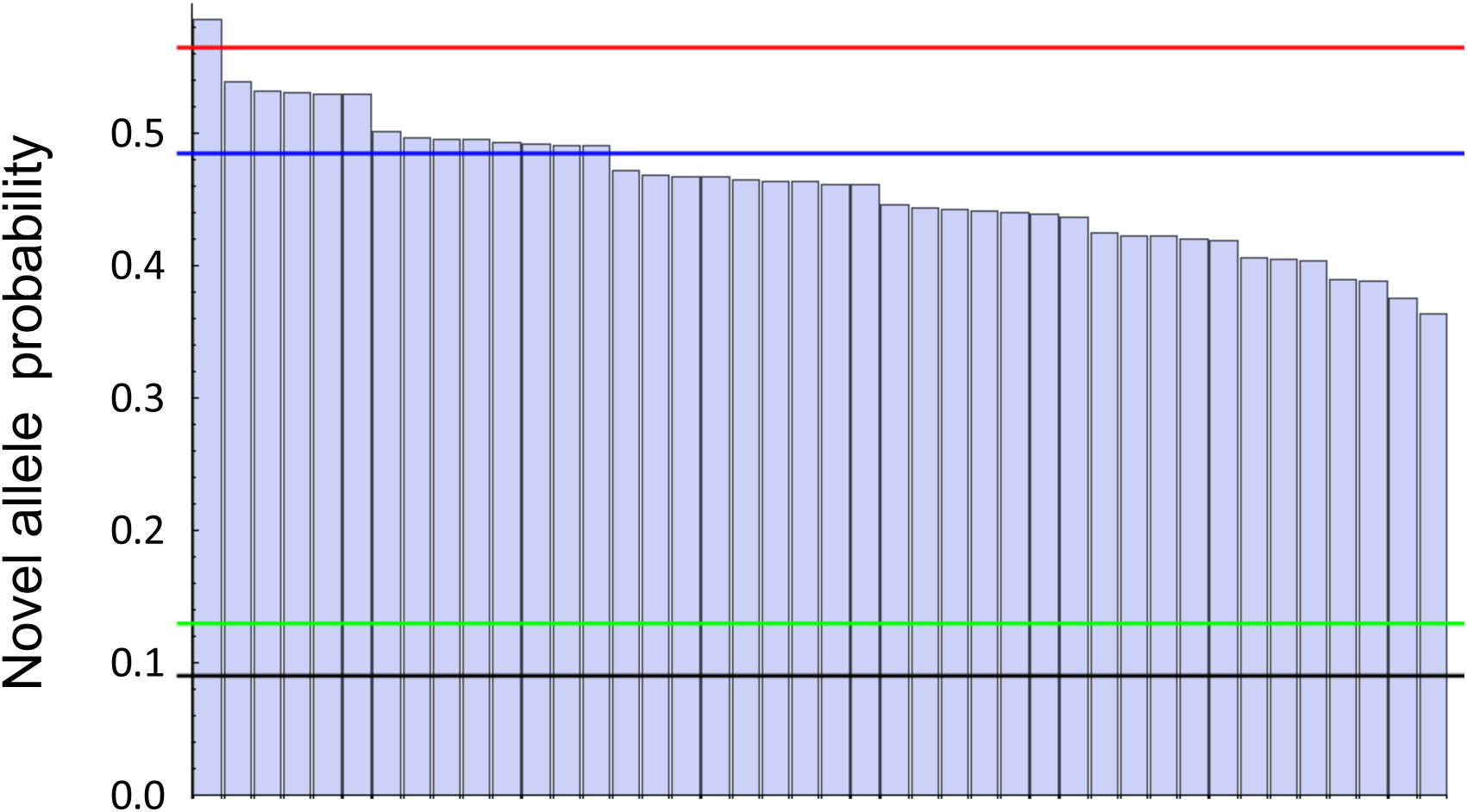
Ranked novel-allele probabilities across AFSs under *θ* = *M*_0_ = *M*_1_ = *c*_0_ = *c*_1_ = 1.0, *n*_0_ = 0, *n*_1_ = 10. Horizontal lines correspond to the marginal novel-allele probability across AFSs (blue (16)), the ESF probability for panmictic populations (black (3a)), the ESF probability with a proposed effective *θ* (green), and the De Iorio and Griffiths (2004b) IS proposal (red (8a)).

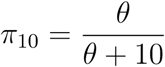

(see (3a)), greatly underestimates the true novel-allele probability for every AFS. The green line, corresponding to the ESF expression with *θ* replaced by an effective *θ*, designed to account in part for population structure (Eq. (26) of Uyenoyama *et al.* 2019), provides only a modest improvement. The IS proposal (8a) of De Iorio and Griffiths (2004b) (red), which takes into account the numbers of lineages in each deme, greatly increases the novel-allele probability, bringing the proposal much closer to the marginal. However, it overestimates the novel-allele probability for all AFSs with the exception of the maximal (leftmost) value, which corresponds to a monomorphic sample.

Figure 3 presents histograms of novel-allele probabilities for an initial sample of size *n*_0_ + *n*_1_ = 10, with the width of the bars proportional to the AFS probabilities prior to the addition of the last (11^*th*^) gene across initial sample configurations (*n*_0_ = 0, 1, …, 10). For a given histogram, the marginal novel-allele probability (16) corresponds to the mean of the histogram. Among the major factors influencing the the probability that the last-sampled gene represents a new allele is the extent of isolation of last gene in the sampling configuration: whether the last gene derives from a deme different from the majority of the sample. Figure 3 indicates that the mode and most of the mass of the distribution of the novel-allele probability tend to increase with greater isolation of the last-sampled gene (smaller values of *n*_0_). This trend is consistent with the observation of Uyenoyama *et al.* (2019, see their Fig. 3) that singleton mutations are more likely to occur on the long branches connecting isolated lineages to the gene genealogy of the rest of the sample. Figure 3 suggests that this effect persists even for mild levels of isolation: sampling configurations in which the last-sampled gene derives from the deme of a minority of the sample, for example. This trend is apparent even for relatively high rates of gene flow (*e.g., M*_0_ = *M*_1_ = 1 in Fig. 3) and intensifies as gene flow declines. Higher values of the scaled mutation parameter tend to increase the novel-allele probability, although the effect depends on the relative magnitudes of the rates of mutation (*θ*), migration (*M*_*i*_), and coalescence (*c*_*i*_).

**Figure 3:**
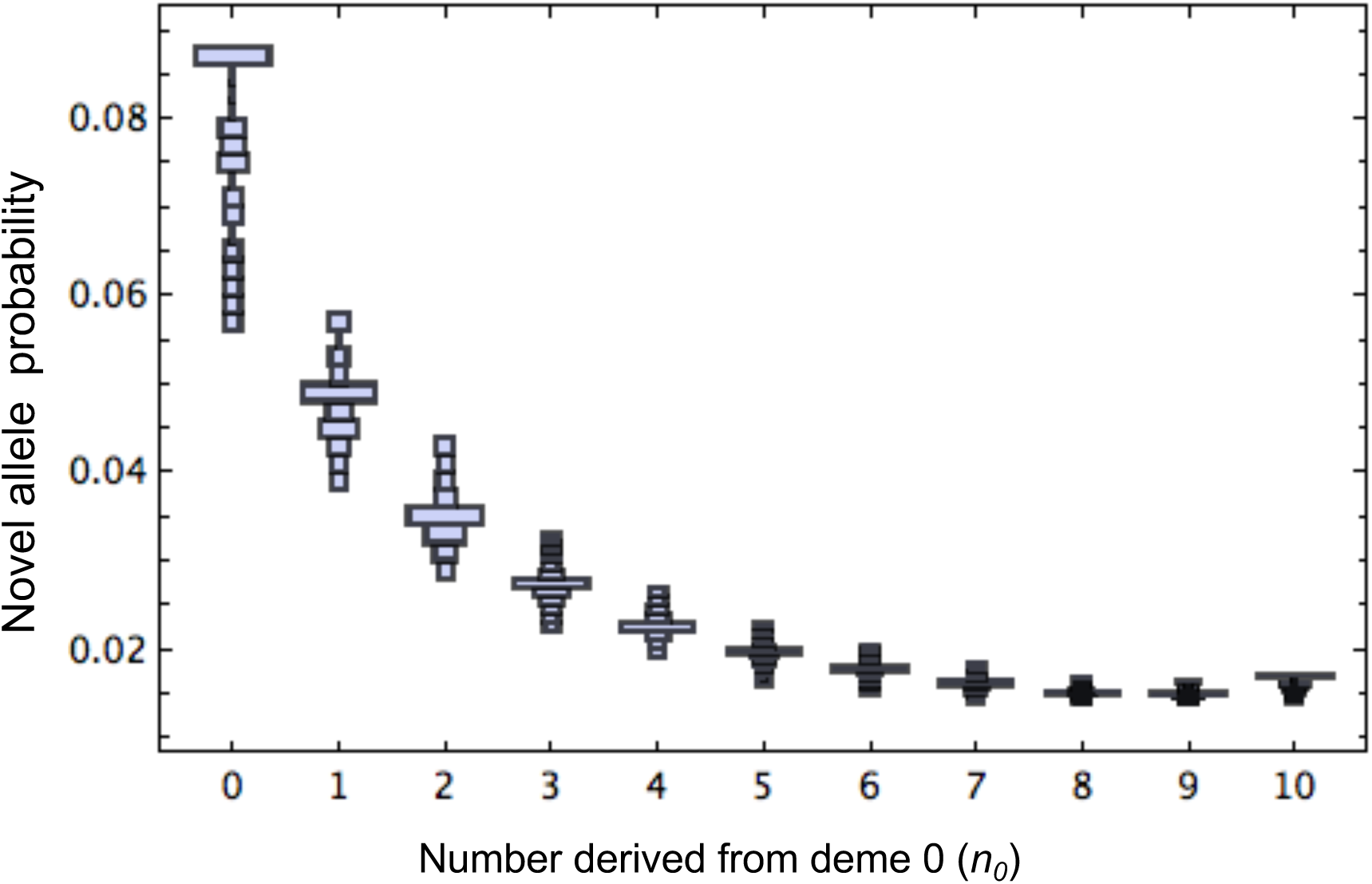
Novel-allele probabilities across AFSs under *θ* = 0.1, *M*_0_ = *M*_1_ = *c*_0_ = *c*_1_ = 1.0, *n*_0_ + *n*_1_ = 10 across initial sample configurations (*n*_0_ = 0, 1, …, 10). All histograms have 10 bins, with bar width proportional to the sum of the probabilities of the AFSs that contribute to each bin.

Under the parameter assignments corresponding to Figure 2, Figure 4 presents the relative error of the IS proposal for the novel-allele probability of De Iorio and Griffiths (2004b), expressed as a relative proportion:

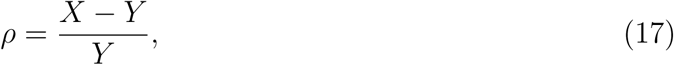

for *X* corresponding to (8a) and Y to the actual marginal novel-allele probability (16). For this symmetric migration case (*M*_0_ = *M*_1_), the IS proposal (8a) shows a general correspondence with the marginal probability of a novel allele (16), with a maximum error of about 17% for the case in which the entire penultimate sample derives from a deme distinct from that of last gene (*n*_0_ = 0, *n*_1_ = 10, as in Fig. 2). Overall, the IS proposal (8a) overestimates the novel-allele probability for samples in which a minority of genes in the original sample derive from the deme from which the last gene is sampled (*n*_0_ ≤ 4). For larger values of *n*_0_, the IS proposal underestimates the novel-allele probability, with the exception of the case in which the entire sample derives from the deme from which the last allele is sampled. Under some conditions (Fig. 3, for example), this case (*n*_0_ = 10) can reverse the trend of declining novel-allele probabilities, but even so, the IS proposal tends to overestimate the marginal novel-allele probability. Figure 5 illustrates a similar qualitative pattern for large sample sizes (*n* = 100), but with higher relative errors.

**Figure 4:**
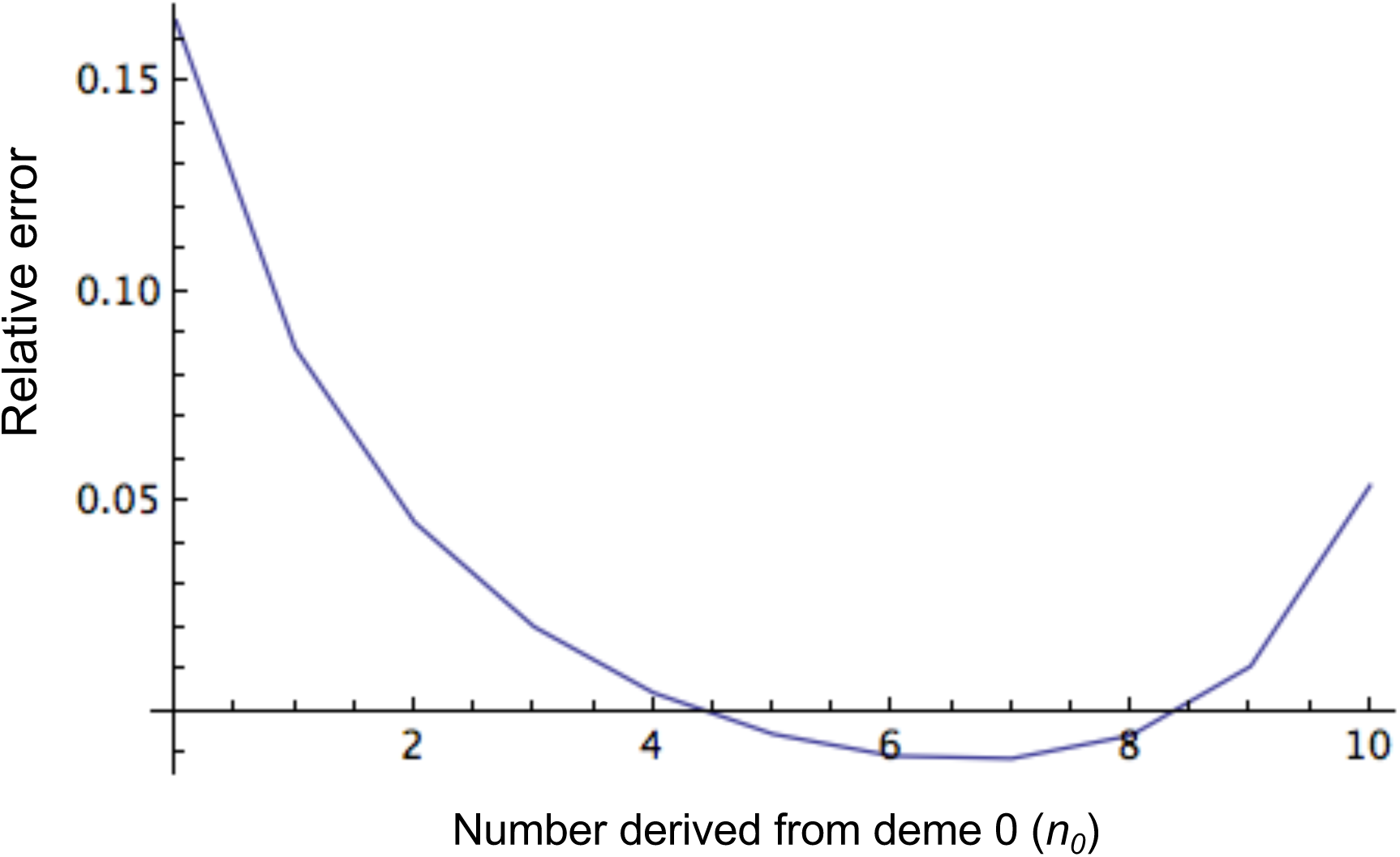
Relative error *ρ* (17) between the marginal probability that a gene sampled from deme 0 represents a novel allele (16) and IS proposal (8a), across initial sampling configurations in which *n*_0_ genes derive from deme 0 (*n*_0_ = 0, 1, …, 10), with *θ* = *M*_0_ = *M*_1_ = *c*_0_ = *c*_1_ = 1.0.

**Figure 5:**
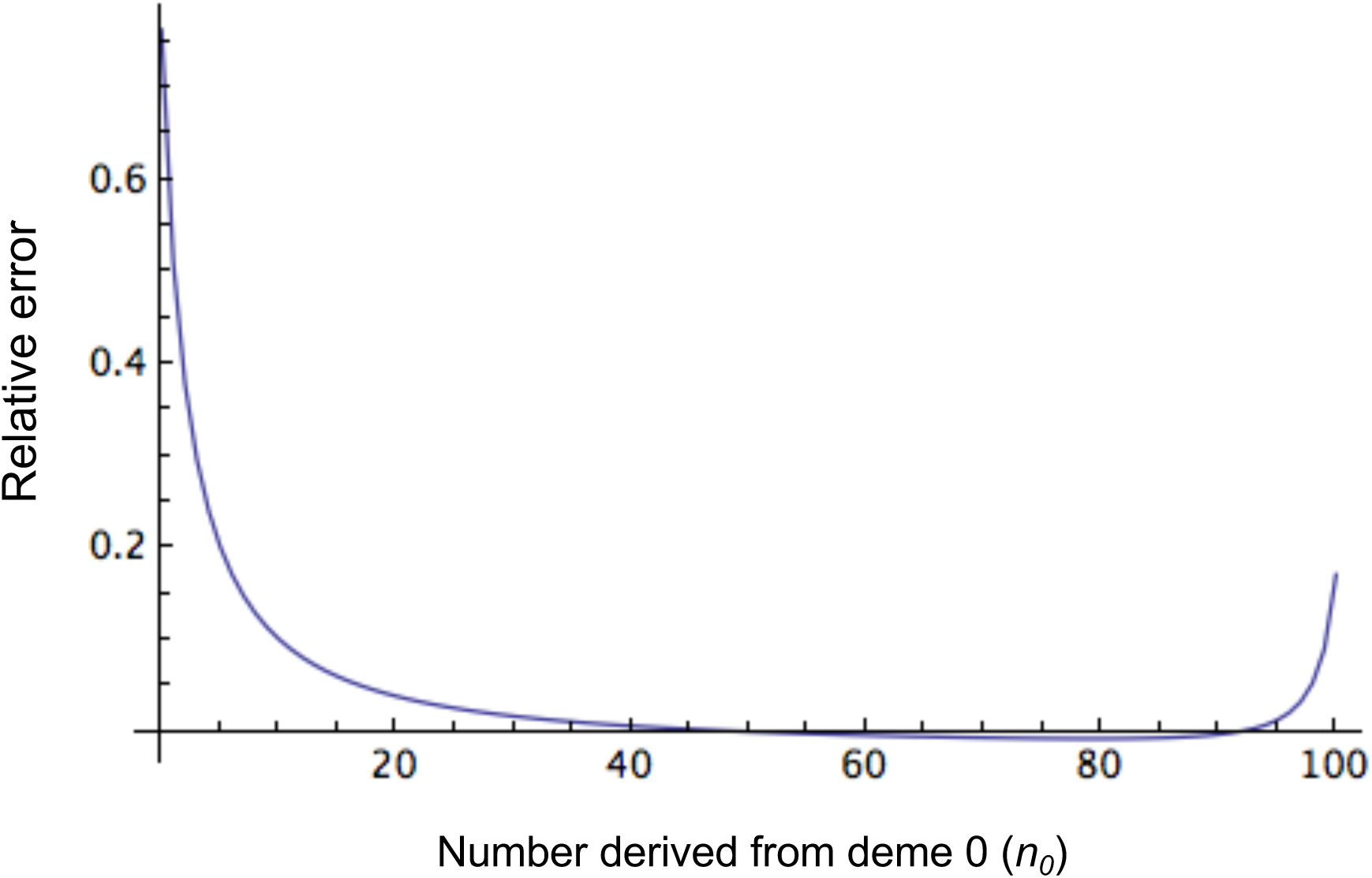
Relative error *ρ* (17) between the marginal probability that a gene sampled from deme 0 represents a novel allele (16) and IS proposal (8a), across numbers of genes in the original sample of size *n* = 100 derived from deme 0 (*n*_0_ = 0, 1, …, 100), with all other parameters as in Fig. 4.

Figure 6 illustrates that the IS proposal (8a) can overestimate by several-fold the marginal novel-allele probability under asymmetric migration (*M*_0_ ≠ *M*_1_), especially in cases in which the entire initial sample derives from the deme (0) from which the last gene is sampled (blue). For sampling configurations in which the minority of the initial sample derives from that deme, the IS proposal (8a) can underestimate the marginal for high rates of backward migration from that deme (*M*_0_). Our preliminary explorations suggest that the absolute magnitude of the relative error diminishes with increases in the scaled mutation rate (*θ*).

**Figure 6:**
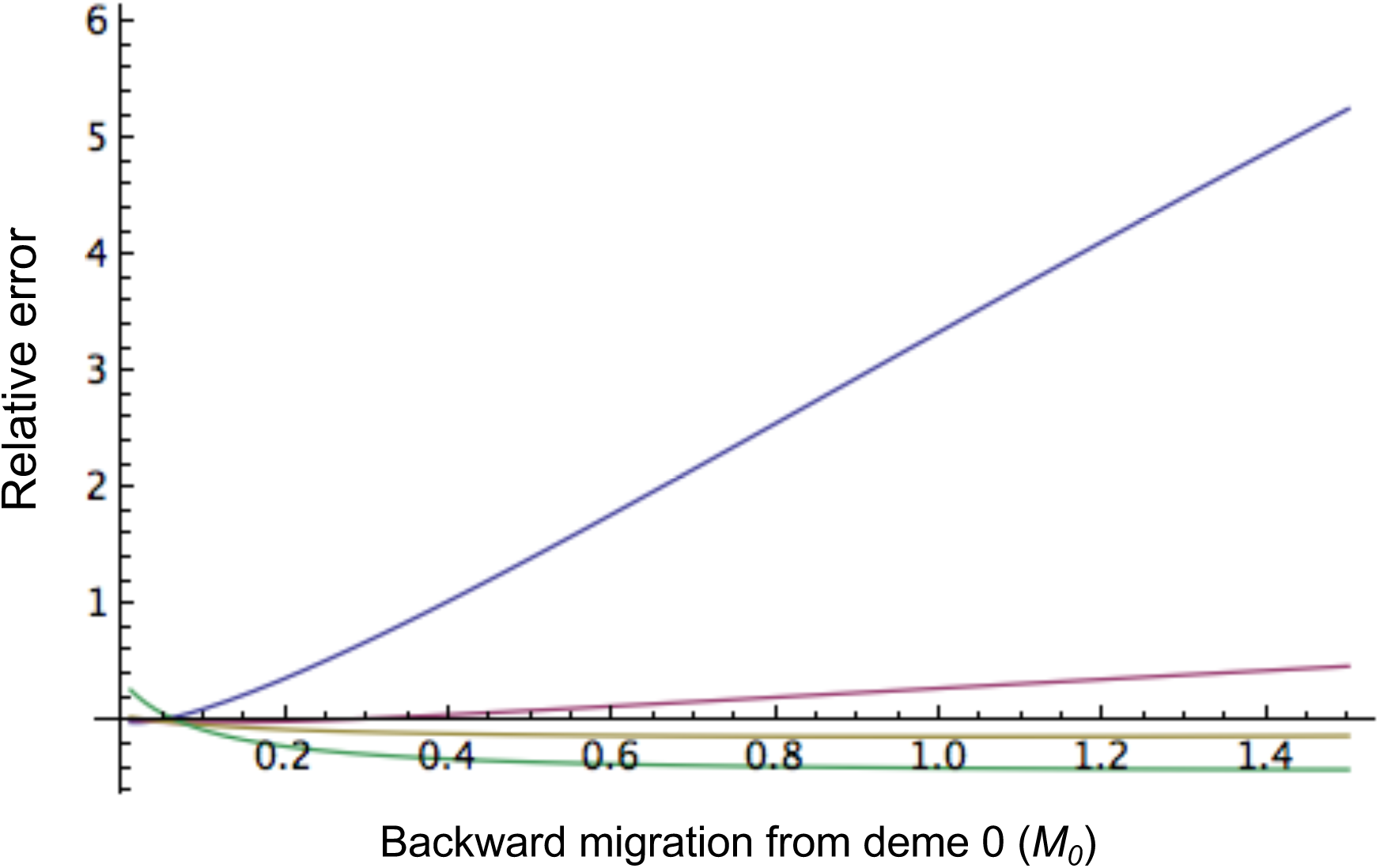
Relative error *ρ* (17) between the marginal probability that a gene sampled from deme 0 represents a novel allele (16) and IS proposal (8a), across rates of backward migration from deme 0 (*M*_0_), with *M*_1_ = 0.05 and *θ* = 0.1 for an initial sample of size *n* = 10. Relative error is highest for initial samples derived entirely from deme 0 (blue, *n*_0_ = 10), declining and becoming negative for samples with progressively more genes derived from deme 1 (magenta, *n*_0_ = 9; yellow, *n*_0_ = 5: green, *n*_0_ = 0).

To explore whether certain characteristics of the penultimate sample (*n* = 10) may provide an indication of the novel-allele probability for the last (11^*th*^) gene, we made pairwise comparisons between the novel-allele probability and other features of an AFS using Kendall’s tau statistic, corrected for ties (Puka 2011). The *i*^*th*^ AFS of the original sample is associated with a novel-allele probability for the last-sampled gene (*x*_*i*_) and also a value (*y*_*i*_) for another feature: for example, *y*_*i*_ may correspond to the number of alleles represented in the *i*^*th*^ AFS. If the novel-allele probability and the other feature were perfectly correlated, then for all pairs of AFSs (*i* and *j*), (*x*_*i*_ − *x*_*j*_) and (*y*_*i*_ − *y*_*j*_) would have the same sign: an increase (decrease) in novel-allele probability is accompanied by an increase (decrease) in the other feature. Of the 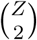 pairwise comparisons among *Z* AFSs, let *C* represent the number of pairs for which the differences are concordant (have the same sign) and *D* the number of discordant pairs (different signs). Let *T*_*x*_ denote the number of pairs of AFSs that show ties for novel-allele probability (*x*_*i*_ = *x*_*j*_) and *T*_*y*_ the number of pairs that show ties for the other feature (*y*_*i*_ = *y*_*j*_). Kendall’s tau-b statistic accommodates such ties:

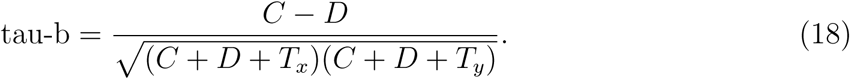

Figure 7 presents this statistic for comparisons between novel-allele probability and other features for an original sample of size *n* = 10 under symmetric migration (*M*_0_ = *M*_1_), with the abscissa (*n*_0_ = 0, 1, …, 10) indicating the number of genes in the original sample derived from deme 0. Interestingly, the concave-down curve (magenta) suggests that novel-allele probability tends to show a negative association with number of alleles in the original sample for samples in which most of the genes derive from a single deme (*n*_0_ close to 0 or *n* = 10). For such unbalanced samples, the next-sampled gene is more likely to be novel if the original sample comprises fewer distinct alleles, whether or not the last gene derives from the deme from which the majority of the original sample derive. However, little association is apparent for balanced samples. The concave-up curve (blue) suggests that the novel-allele probability is generally higher for AFSs that themselves have higher probability, a trend that persists under higher scaled mutation rates (Fig. 8). The remaining curve suggests that higher proportions of private alleles (those observed in the subsample derived from exactly one deme; see Slatkin 1985) tend to be positively associated with higher novel allele probabilities. Figure 8 suggests that the associations may be strengthened under a higher mutation rate (*θ* = 1.0).

**Figure 7:**
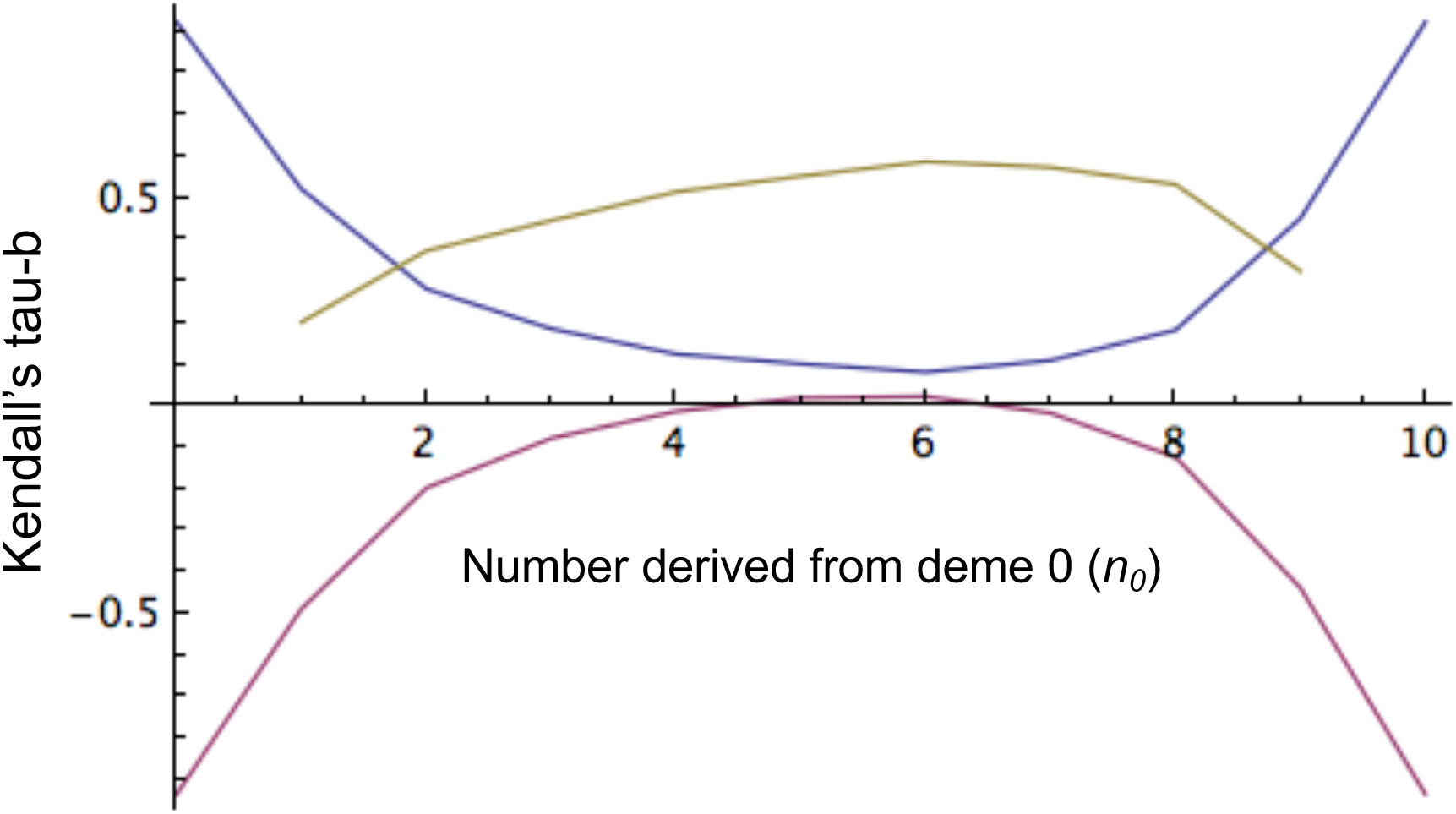
Kendall’s tau-b (18) measure of association across AFSs between the probability that an additional gene sampled from deme 0 represents a novel allele for a given AFS and another feature of the AFS for an original sample of size 10 with *n*_0_ genes (*n*_0_ = 0, 1, …, 10) derived from deme 0, with *c*_0_ = *c*_1_ = *M*_0_ = *M*_1_ = 1 and *θ* = 0.1. The concave-up curve (blue) corresponds the the probability of the original AFS, the concave-down curve (magenta) to the number of alleles observed in the original sample, and the remaining curve to the proportion of alleles observed in only a single deme.

**Figure 8:**
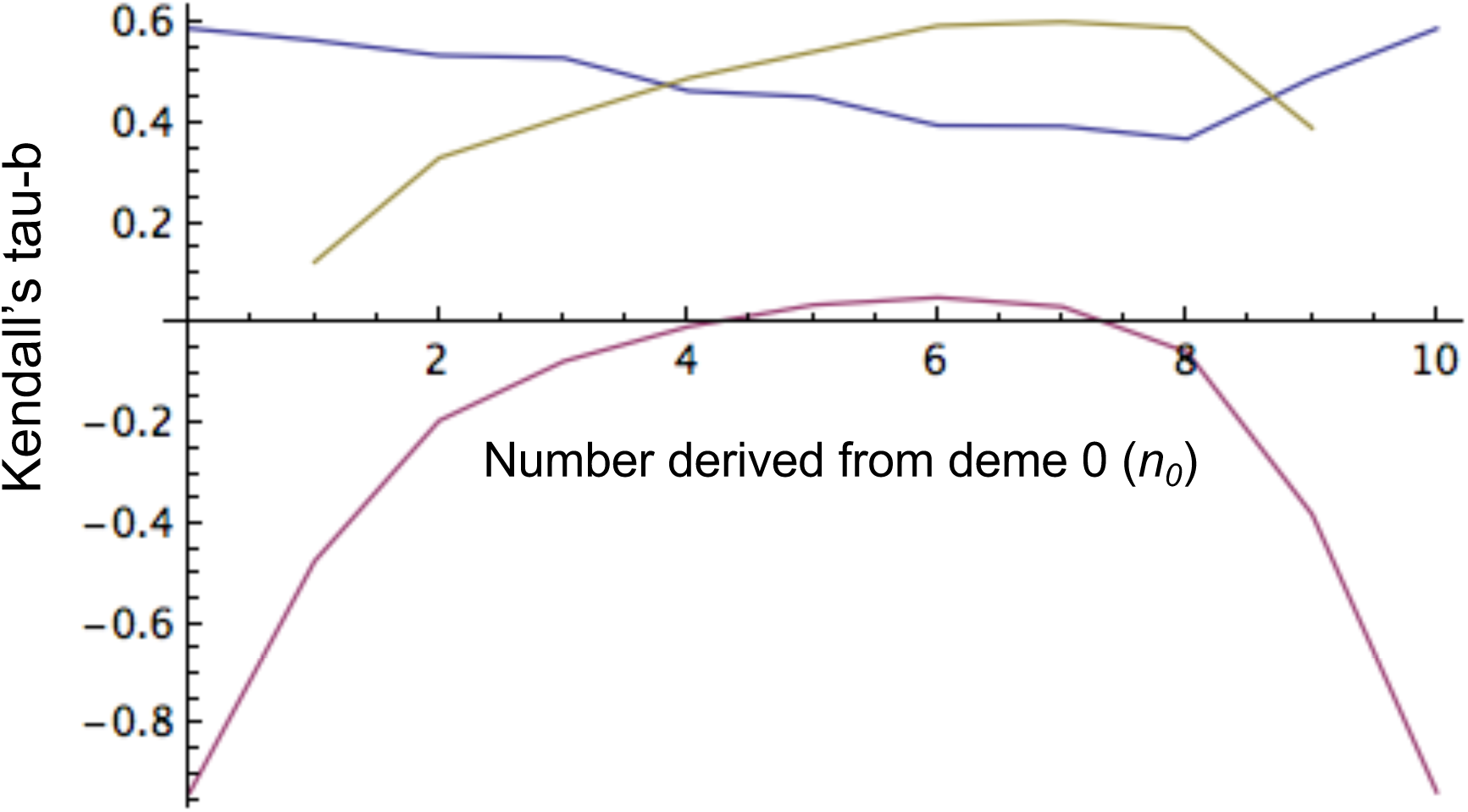
Kendall’s tau-b (18) as described for Fig. 7, but with *θ* = 1.0.

## 5 Discussion

An alternative to the separate proposal of an unlabelled genealogy and then a mutational history conditional on the genealogy entails the proposal of full labelled histories. Our inductive method (Uyenoyama *et al.* 2019) uses the labelled coalescence argument of Karlin and McGregor (1972) to determine the probability of all allele frequency spectra (AFSs) in structured populations under the infinite-alleles model of mutation. Because the number of AFSs grows rapidly (although slower than exponentially) with sample size (*n*), at the approximate rate

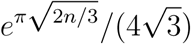

(Bóna 2011, p. 98), the computation of all AFS probabilities is clearly impractical for large samples. For samples comprising 100 genes derived entirely from a single deme, 190,569,292 distinct AFSs exist (Abramowitz and Stegun 1965, Table 21.5), and the total number, over all possible sampling configurations, is many orders of magnitude greater. Even so, our recursive method (16) can determine the exact marginal novel-allele probability for large samples, including *n* = 100 (Fig. 5). A similar recursion (Eqn. (14) of Uyenoyama *et al.* 2019) provides the probability generating function of the total number of alleles observed in samples of a given size.

De Iorio and Griffiths and colleagues (see especially De Iorio and Griffiths 2004a,b; De Iorio *et al.* 2005) have developed a class of importance sampling (IS) proposals for the determination of likelihoods that can accommodate generalized mutation and migration models in structured populations. Those IS proposals, which appear to be the most efficient available, were constructed by extrapolating fundamental properties of the ESF to structured populations (see Section 3.1). Key to this approach is their approximation of the probability, given an arbitrary sample AFS, that the next-observed gene represents a novel allelic class. Our study of this approximation in the simple model explored here (6) suggests that the success of their IS proposals may reflect in part the similarity of their novel-allele probability (8a) to the true marginal novel-allele probability (16) across AFSs of the sample prior to the addition of the last-observed gene.

We find that the IS proposal for the novel-allele probability (8a) tends to overestimate the marginal (16) for cases in which the last-observed gene is drawn from a deme different from that from which the majority of the sample was derived (Figs. 4 and 5). In such cases, relatively few of the existing lineages reside in the same deme as the lineage of the last-observed gene. A more ancient coalescence of the lineage of the last-observed gene allows more time for a mutation to occur in that lineage, tending to increase the novel-allele probability. While the novel-allele probability is indeed relatively high for such sampling configurations (*e.g.*, see Fig. 3), the IS proposal (8) favors even higher values. We suggest that substituting the actual marginal novel-allele probability (16) for (8) and otherwise preserving the De Iorio and Griffiths (2004b) method for constructing IS proposals may improve efficiency.

Our preliminary explorations of the marginal probability of a novel allele (16) suggest some additional qualitative trends. Most strikingly, genes derived from a deme different from that of the majority of the sample have higher probabilities of representing a novel allelic class (Fig. 3). For unbalanced samples (unequal numbers of genes derived from the demes), the novel-allele probability tends to show a negative association with the number of alleles in sample (Figs. 7 and 8): the novel-allele probability is higher for initial samples comprising fewer alleles. The distribution of allele number (section 2 of Uyenoyama *et al.* 2019) can be used to ascertain whether the number of alleles in the initial sample is low. In addition, observation in the initial sample of more private alleles (those observed in the subsample derived from a single deme) appears to be associated with higher novel-allele probabilities. Observation of private alleles may suggest low migration rates (see Slatkin 1985), again allowing more time for a mutation to occur in the lineage of the last-observed gene.

## Acknowledgments

We thank Editor Noah Rosenberg and the Editorial Board for the opportunity to contribute to this volume, celebrating the 50^*th*^ anniversary of *Theoretical Population Biology*, the forum in which the Ewens Sampling Formula (Ewens 1972; Karlin and McGregor 1972) and many other works on which we have drawn have appeared. We are grateful to the Editor and the reviewers for their helpful comments. Public Health Service grant GM 37841 (MKU) provided partial funding for this research.

## Appendix A ESF recursion

In using the labelled coalescence argument (4) to derive the ESF, Karlin and McGregor (1972) conditioned on two events: descent of the *n* genes in the present sample from *n* − 1 distinct genes in the preceding generation, with probability

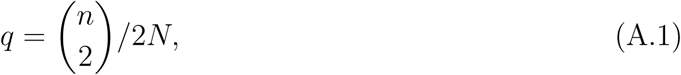

and descent from *n* distinct genes with the complement probability (1 − *q*). Under the large *N* assumption, all other possible derivations of the sample occur at negligible rates.

For monomorphic samples, which comprise *n* copies of a single allele (*a*_*n*_ = 1),

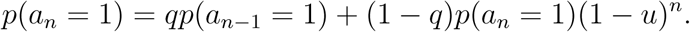

Substituting (A.1) and ignoring terms of second order or smaller in rates of coalescence (1*/*2*N*) or mutation (*u*) yields a familiar expression:

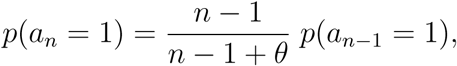

in which the (*n* − 1)*/*(*n* − 1 + *θ*) term represents the probability that the most recent event (*T*) corresponds to the coalescence of a pair of lineages. Substitution of (1) verifies the ESF in this case.

For samples comprising more than a single allele (*a*_*n*_ = 0), the recursion corresponds to

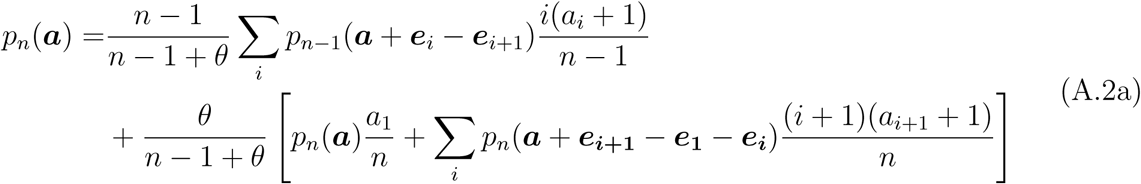

in which ***e***_***i***_ denotes a unit vector, with unity in the *i*^*th*^ position and zeros elsewhere and unmeaningful expressions (*e.g.*, probability of spectra with negative elements) are defined as zero (Karlin and McGregor 1972). The first term on the right of (A.2a) corresponds to the splitting of an allelic lineage (in an ancestral sample of size *n* − 1) as the most recent evolutionary event (*T*) and the bracketed term to mutation (in an ancestral sample of size *n*). Mutation in a singleton allele preserves the AFS ***a***, mutation in an allele represented exactly twice generates two additional singletons, and mutation in an allele represented *i* times (*i* > 2) generates one singleton and reduces the multiplicity of the allele to *i* − 1. Rearrangement of (A.2a) produces

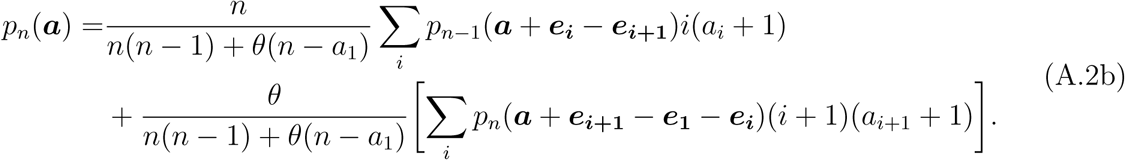

Tavaré (2004, his equation (3.5.2)) has derived an identical equation. Substitution of (1) verifies the ESF as a solution.

## Appendix B Probability that the last-sampled gene represents a novel allele

Here, we use the conditional probability interpretation of ratios of AFS probabilities (5) noted by Karlin and McGregor (1972) to provide an alternative derivation of the probability (3a) that the next-sampled (*n*^*th*^) gene represents a novel allele:

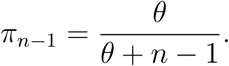

Dividing both sides of fundamental recursion (A.2b) by *p*_*n*_(***a***), we obtain:

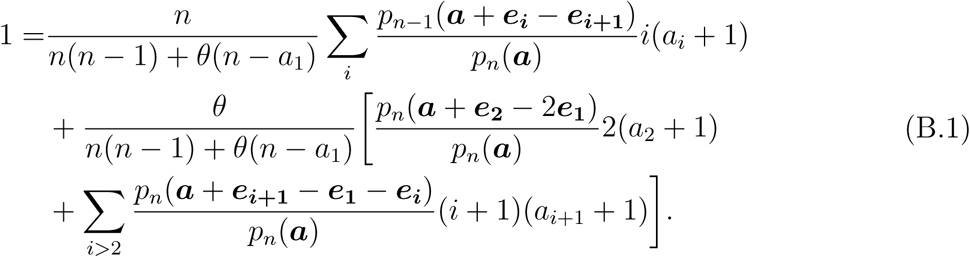

We denote the probability that the last-sampled gene represents an allele that occurs in the full sample with multiplicity *i* by

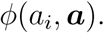

For any non-singleton in the full sample,

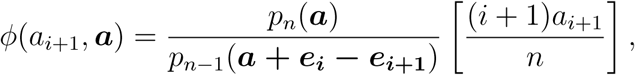

in which the second factor reflects the probability that one of the genes representing an allele with multiplicity *i* + 1 in the full sample is sampled last (compare (5)). An alternative expression conditions on the last-sampled gene representing an allelic type already observed among the first *n* − 1 genes:

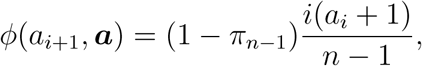

in which the second factor reflects that the allelic class of the last-sampled gene corresponds to the class of a gene sampled uniformly at random from the sample at size *n* − 1. Equating these expressions for *ϕ*(*a*_*i*+1_, ***a***) yields

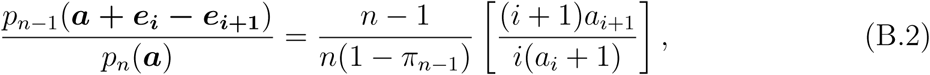

the ratio of AFS probabilities in the first summation of (B.1).

The second ratio of AFS probabilities in (B.1) corresponds to a product of conditional probabilities:

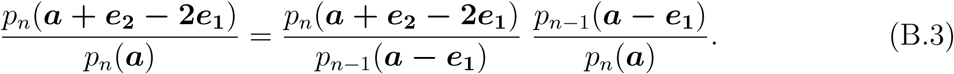

The probability that the last-sampled gene represents a doubleton allele in a full sample with AFS ***a* + *e***_**2**_ ***−* 2*e***_**1**_ is

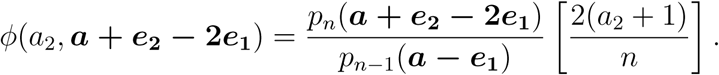

This expression is also equal to the probability that the last-sampled gene is not novel relative to the penultimate sample (1 − *π*_*n*−1_) and belongs to an allelic class already represented by a singleton:

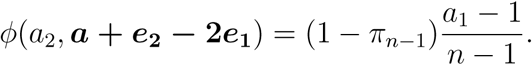

Equating these expressions yields

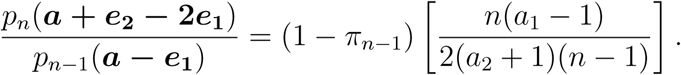

Substitution of this expression and (5) into (B.3) produces

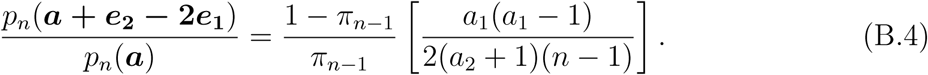

The final ratio of AFS probabilities in (B.1) corresponds to

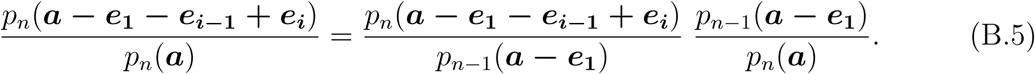

Once again, we have two expressions for the probability that the last-sampled gene represents an allele with multiplicity *i* in a sample of size *n* with AFS ***a − e***_**1**_ ***− e***_***i−*1**_ **+ *e***_***i***_:

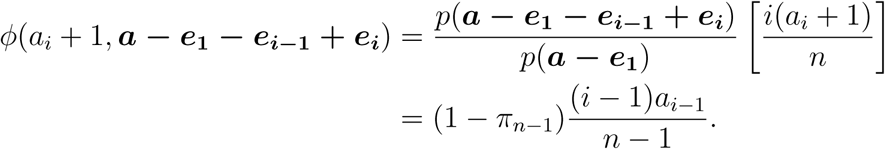

Together with (5), these expressions produce

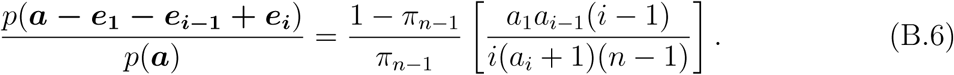

Substitution of (B.2), (B.4), and (B.6) into (B.1) produces a quadratic in *π*_*n*−1_:

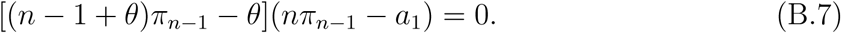

Ewens’s (1972) expression (3a) for the probability of sampling novel allele on the *n*^*th*^ draw is indeed a root of this equation. The second root, independent of the scaled mutation parameter *θ*, simply represents the probability that the last-sampled gene in a sample of size *n* is one of the *a*_1_ singletons (*a*_1_*/n*). As it is clear that the novel-allele probability must depend on *θ*, the first root is this probability.

## Appendix C Proposing an ancestor given the descendant under the ESF

The most efficient IS proposal for generating the genealogical history of the sample would incorporate the actual distribution of ancestor *A* in which the most recent evolutionary event occurs, given descendant *D*:

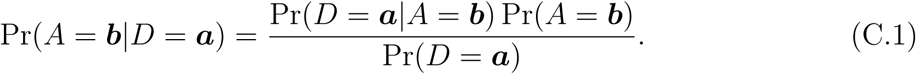

While determining the conditional probability of descendant *D* given ancestor *A* is straight-forward, this expression illustrates that determination of the reverse conditional probability is tantamount to full solution of the likelihood recursion (Stephens and Donnelly 2000).

Observation of *D* excludes certain AFSs from consideration as *A*: for example, AFSs that require more than one evolutionary event to be transformed into *D* have Pr(*D*|*A*) = 0. However, the conditional distribution of *A* is not uniform over non-excluded ancestral states. To explore a means of proposing *A* from *D*, Hobolth *et al.* (2008) examined the conditional distribution (C.1) under the ESF (1):

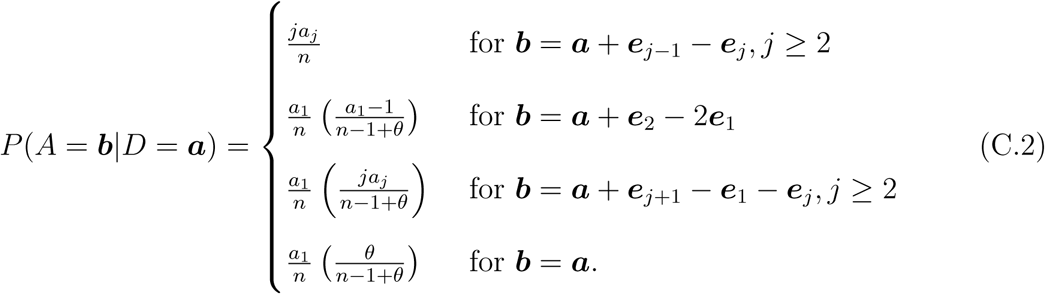

These expressions suggest choosing a gene uniformly at random from the sample as the lineage that participated in the most recent event. This gene either occurs in the sample with multiplicity greater than or equal to 2 or is a singleton. With probability *ja*_*j*_*/n*, the chosen gene represents an allele that occurs in the sample in multiplicity *j* (*j* ≥ 2). In this case, the most recent event must have been a coalescence between that lineage and another representative of the same allelic class, which implies ***b*** = ***a*** + ***e***_*j*−1_ − ***e***_*j*_.

Alternatively, with probability *a*_1_*/n*, the focal gene represents a singleton allele, newly-arisen by mutation. Using that all sampling orders are equiprobable, we regard the focal gene as the last-sampled gene. Immediately ancestral to the mutational event that created the allelic class of the focal gene, the focal lineage represented a singleton allele with probability *θ/*(*n* − 1 + *θ*). In this case, the state of the ancestor was ***b*** = ***a***. Otherwise, with probability (*n* − 1)*/*(*n* − 1 + *θ*), the focal lineage shared its allelic class with at least one of the other *n* − 1 lineages. To determine the allelic class of the focal lineage, we choose a gene uniformly at random from the other *n* − 1 lineages and assume the focal gene shares its allelic class. A singleton allele relative to the *n* − 1 non-focal lineages is chosen with probability (*a*_1_ − 1)*/*(*n* − 1), implying that the focal gene represented a doubleton allele immediately ancestral to the most recent event (mutation). Accordingly, *A* = ***a*** + ***e***_2_ − 2***e***_1_ with probability

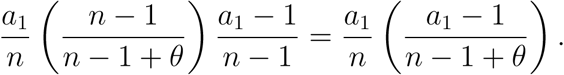

With probability *ja*_*j*_*/*(*n* − 1), the gene chosen from the *n* − 1 non-focal lineages represents an allele with multiplicity *j* ≥ 2, which implies that *A* = ***a*** + ***e***_*j*+1_ − ***e***_1_ − ***e***_*j*_ with probability

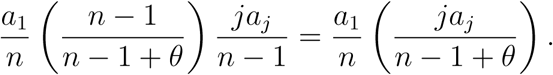

## Figure captions

**Figure 1**. Topology consistent with observation of fixed mutational differences between sub-samples derived from each of two species (red and blue). Mutations arising on the branch labelled *f/a* are fixed (*f*) in the subsample derived from the red species and absent (*a*) from the subsample derived from the blue species. Similarly, mutations arising on the branch labelled *a/f* occur only in the subsample derived from the blue species.

**Figure 2**. Ranked novel-allele probabilities across AFSs under *θ* = 0.5, *M*_0_ = *M*_1_ = 0.1, *c*_0_ = *c*_1_ = 1, *n*_0_ = 0, *n*_1_ = 10. Horizontal lines correspond to the marginal novel-allele probability across AFSs (blue (16)), the ESF probability for panmictic populations (black (3a)), the ESF probability with a proposed effective *θ* (green), and the De Iorio and Griffiths (2004b) IS proposal (red (8a)).

**Figure 3**. Novel-allele probabilities across AFSs under *θ* = 0.1, *M*_0_ = *M*_1_ = *c*_0_ = *c*_1_ = 1.0, *n*_0_ + *n*_1_ = 10 across initial sample configurations (*n*_0_ = 0, 1, …, 10). All histograms have 10 bins, with bar width proportional to the sum of the probabilities of the AFSs that contribute to each bin.

**Figure 4**. Relative error *ρ* (17) between the marginal probability that a gene sampled from deme 0 represents a novel allele (16) and IS proposal (8a), across initial sampling configurations in which *n*_0_ genes derive from deme 0 (*n*_0_ = 0, 1, …, 10), with *θ* = 0.5, *M*_0_ = *M*_1_ = 0.1, and *c*_0_ = *c*_1_ = 1.

**Figure 5**. Relative error *ρ* (17) between the marginal probability that a gene sampled from deme 0 represents a novel allele (16) and IS proposal (8a), across numbers of genes in the original sample of size *n* = 100 derived from deme 0, for symmetric backward migration rates *M*_0_ = *M*_1_ = 0.5 and *θ* = 0.5.

**Figure 6**. Relative error *ρ* (17) between the marginal probability that a gene sampled from deme 0 represents a novel allele (16) and IS proposal (8a), across rates of backward migration from deme 0 (*M*_0_), with *M*_1_ = 0.05 and *θ* = 0.1 for an initial sample of size *n* = 10. Relative error is highest for initial samples derived entirely from deme 0 (blue), declining and becoming negative for samples with progressively more genes derived from deme 1 (other curves).

**Figure 7**. Kendall’s tau-b (18) measure of association across AFSs between the probability that an additional gene sampled from deme 0 represents a novel allele for a given AFS and another feature of the AFS for an original sample of size 10 with *n*_0_ genes (*n*_0_ = 0, 1, …, 10) derived from deme 0, with *c*_0_ = *c*_1_ = *M*_0_ = *M*_1_ = 1 and *θ* = 0.1. The concave-up curve (blue) corresponds the the probability of the original AFS, the concave-down curve (magenta) to the number of alleles observed in the original sample, and the remaining curve to the proportion of alleles observed in only a single deme.

**Figure 8**. Kendall’s tau-b (18) as described for Fig. 7, but with *θ* = 1.0.

